# Major changes in domain arrangements are associated with the evolution of termite castes

**DOI:** 10.1101/2023.05.15.540413

**Authors:** Alina A. Mikhailova, Elias Dohmen, Mark C. Harrison

## Abstract

Domains as functional protein units and their rearrangements along the phylogeny can shed light on the functional changes of proteomes associated with the evolution of complex traits like eusociality. This complex trait is associated with sterile soldiers and workers, and long-lived, highly fecund reproductives. Unlike in Hymenotpera (ants, bees, and wasps), the evolution of eusociality within Blattodea, where termites evolved from within cockroaches, was accompanied by a reduction in proteome size, raising the question of whether functional novelty was achieved with existing rather than novel proteins. To address this, we investigated the role of domain rearrangements during the evolution of termite eusociality. Analysing domain rearrangements in the proteomes of three solitary cockroaches and five eusocial termites, we inferred more than 5000 rearrangements over the phylogeny of Blattodea. The 90 novel domain arrangements that emerged at the origin of termites were enriched for several functions related to longevity, such as protein homeostasis, DNA repair, mitochondrial activity, and nutrient sensing. Many domain rearrangements were related to changes in developmental pathways, important for the emergence of novel castes. Along with the elaboration of social complexity, including permanently sterile workers and larger, foraging colonies, we found 110 further domain arrangements with functions related to protein glycosylation and ion transport. We found an enrichment of caste-biased expression and splicing within rearranged genes, highlighting their importance for the evolution of castes. Furthermore, we found increased levels of DNA methylation among rearranged compared to non-rearranged genes suggesting fundamental differences in their regulation. Our findings indicate an importance of domain rearrangements in the generation of functional novelty necessary for termite eusociality to evolve.

## Introduction

Protein domains are the functional and structural units of proteins (Forslund and Sonnhammer, 2012; Dohmen et al., 2020). Domains can be gained by a protein, or the relative positions of existing domains to each other within a domain arrangement or domain architecture (DA) can be rearranged, allowing a protein to acquire novel functions (Moore et al., 2008). These rearrangements can occur via different mechanisms, including gene fusion, exon shuffling, recombination and gene duplication, and are important for the emergence of various traits (Patthy, 2003; Moore et al., 2008; Buljan et al., 2010; Forslund et al., 2019). Particularly, the evolution of multicellularity, one of the major evolutionary transitions (Szathmáry and Smith, 1995), was accompanied by an increase in domain rearrangements (Ekman et al., 2007). This is demonstrated, for example, in extracellular communication, extracellular matrix formation and intracellular signalling pathways, in which novel multi-domain proteins played an important evolutionary role (Patthy, 1985; Pawson, 1995; Patthy, 2003), as well as in the evolution of blood coagulation cascade proteins in vertebrates (Coban et al., 2022).

Not only the reuse of existing domains can be a source of evolutionary innovation, but also the emergence of novel domains can lead to the evolution of new traits. It was shown that the rapid emergence of animal-specific domains contributed to the functional diversification in animals as well as the evolution of tyrosine phosphorylation systems and coagulation cascades (Itoh et al., 2007). Also, a high rate of domain emergences was previously observed at the root of mammals and was potentially linked to the emergence of hair and the evolution of the mammalian immune system (Dohmen et al., 2020). Analysing types and rates of domain rearrangements and emergences along a phylogeny can help us to detect molecular events leading to evolutionary innovation. For example, in arthropods 40 000 domain rearrangements could be identified, contributing to the enormous diversity that exists especially within insects (Thomas et al., 2020).

The role of domain rearrangements in the next major transition, from multicellular individuals to eusocial colonies (Szathmáry and Smith, 1995), has so far received little attention. This complex trait, which is characterized by cooperative brood care, overlapping generations and division of labour (Wilson and Hölldobler, 2005), has emerged in such divergent orders as Coleoptera (ambrosia beetles), Hymenoptera (bees, wasps and ants) and Blattodea, where termites have evolved within the cockroaches (Kent and Simpson, 1992; Crespi, 1996; Inward et al., 2007; Lo et al., 2007; Sherman et al., 2017). It has been hypothesised that, while mainly regulatory changes are expected during the early stages of social evolution, greater functional changes occur with the evolution of greater social complexity along with the emergence of distinct castes (Rehan and Toth, 2015). These expectations have been largely confirmed by comparative genomic analyses on the transition to eusociality in bees (Kapheim et al., 2015; Shell et al., 2021), ants (Simola et al., 2013) and termites (Harrison et al., 2018) such as increased transcriptional regulation and evolution of gene families involved in chemoperception. The potential role of domain evolution in this major transition has been highlighted by novel DAs being found at the node representing one origin of eusociality in bees, affecting proteins with functions related to embryonic morphogenesis, cell growth and insulin signalling (Dohmen et al., 2020).

Termites represent an interesting clade for investigating the molecular evolution of eusociality, since extant species display varying levels of social complexity, allowing the inference and mapping of various social traits to the phylogeny (Korb and Thorne, 2017). Furthermore, as with most eusocial taxa, reproductive individuals display extreme lifespans despite high fertility. In the higher termites, of the genus *Macrotermes*, for example, queens can live for over 20 years (Elsner et al., 2018) while producing thousands of eggs per day (Kaib et al., 2001). This convergent decoupling of the longevity/fecundity trade-off in eusocial orgnaisms has been linked to an upregulation of members of the Insulin/insulin-like growth factor (IGF-1) signaling (IIS) pathway, as well as increased expression of mitochondrial and DNA repair functions (Séité et al., 2022; Monroy Kuhn et al., 2019). Sterile soldiers emerged once early in the evolution of termites and are present in all extant lineages except for a few Termitidae genera where soldiers were secondarily lost (Noirot and Pasteels, 1987). The origin of the sterile worker caste that consists of morphologically specialized individuals which diverge early from the imaginal (reproductive) line is considered polyphyletic and correlates with nesting and feeding behaviour (Noirot and Pasteels, 1987; Noirot, 1988; Legendre et al., 2008; Roisin and Korb, 2011). Wood-dwelling termites live in small, non-foraging colonies with totipotent workers, which retain the potential to reach sexual maturity (Korb and Hartfelder, 2008). In the larger, foraging colonies of Mastotermitidae, Hodotermitidae and Termitidae, as well as most Rhinotermitidae species, on the other hand, development follows an early bifurcation leading to the production of true workers without the ability to reach sexual maturation (Abe, 1990; Roisin and Korb, 2011; Pull and McMahon, 2020). The ability of workers to switch from the sterile, apterous to the nymphal-alate line to obtain a certain level of fertility remains to varying degrees in these groups but is generally rare (Roisin and Korb, 2011). In contrast to findings for the evolution of eusociality in Hymenoptera (Kapheim et al., 2015; Shell et al., 2021; Simola et al., 2013), a comparative genomic study into the origins of blattodean eusociality found the unexpected pattern of reductions in proteome size due to gene family contractions in termites compared to non-social outgroups (Harrison et al., 2018). We hypothesise, therefore, that changes in existing proteins, for example, via domain rearrangements, may have played a role in the emergence and elaboration of eusociality in termites.

In this study, we tested this hypothesis by analysing the domain content of available genomes from eight blattodean species with varying levels of social complexity. Beside three, non-social cockroaches (*Blattella germanica, Periplaneta americana* and *Diploptera punctata*), we analysed two lower, wood-dwelling termites (*Zootermopsis nevadensis* and *Cryptotermes secundus*) that form small colonies with totipotent workers consisting of around 900 (Abe, 1990) and 300-400 (Korb and Lenz, 2004) individuals, respectively, in which reproductives live up to around 7 years (Monroy Kuhn et al., 2019; Thorne et al., 2002). To represent termites with higher social complexity, we included three foraging termite species with sterile (‘true’) workers (*Coptotermes formosanus, Reticulitermes speratus*, and *Macrotermes natalensis*) that form large, complex colonies containing up to 50,000 (*C. formosanus*) and 200,000 individuals (Abe, 1990; Meyer et al., 2000), in which reproductives have even longer lifespans of over 10 years (Monroy Kuhn et al., 2019; Keller, 1998). We inferred domain rearrangements that occurred at the origin of termites and at the evolution of sterile workers to understand changes in protein content related to the emergence and elaboration of termite eusociality.

## Methods

### Domain rearrangements calculation

We used previously published proteomes of three cockroaches (*B. germanica, D. punctata, P. americana*), five termites (*Z. nevadensis, C*.*secundus, R. speratus, C. formosanus, M. natalensis*) and two outgroup species (*Eriosoma lanigerum, Frankliniella occidentalis*) for this study (Harrison et al., 2018; Li et al., 2018; Fouks et al., 2022; Terrapon et al., 2014; Shigenobu et al., 2022; Itakura et al., 2020; Poulsen et al., 2014; Biello et al., 2021; Rotenberg et al., 2020). All proteomes filtered for possible pseudogenes and only longest isoforms were kept, using DW-Helper scripts (https://domain-world.zivgitlabpages.uni-muenster.de/dw-helper/index.html). Filtered proteomes were then annotated with Pfam domains in version 30.0 (Mistry et al., 2021) with PfamScan v1.6 (Finn et al., 2016). Completeness of the annotated proteomes was assessed with DOGMA (Dohmen et al., 2016; Kemena et al., 2019) (Supplementary Table 1). A species tree was constructed with OrthoFinder using single-copy orthologs (Emms and Kelly, 2019). DomRates was run on the species tree and annotated proteomes to reconstruct domain rearrangements along the phylogeny (Dohmen et al., 2020). Only exact solutions of DomRates were used in the downstream analyses.

### Gene Ontology enrichment analysis

To infer functional adaptation along the evolution of termites, we performed GO-term enrichment analyses of (i) domain architectures (DA) and (ii) genes with rearranged DAs. For DAs, we used pfam2go to map GO-terms to the corresponding domains in the DAs (Mitchell et al., 2015). The GO universe for this analysis contained all domain rearrangements in Blattodea, and we took domain rearrangements on each inner node of the Blattodea phylogeny as the test set.

To analyse the functions of genes with rearranged domains, we extracted genes containing DAs from the DomRates output for the nodes of the origin of termites (*Z. nevadensis, C. secundus, R. speratus, M. natalensis* species) and the origin of true workers (*R. speratus* and *M. natalensis* species). For each species, we looked at the enriched GO-terms in the genes with domains that were rearranged in one of the two origins.

Both GO-term enrichment analyses were performed with the package topGO in R using the classic and weight algorithms (Alexa et al., 2010). Significantly enriched GO-terms (p-value *<* 0.05) were visualized using the library tagcloud in R (https://cran.r-project.org/web/packages/tagcloud/).

### Differential gene expression analyses

To look into the enrichment of caste-biased genes among genes rearranged at the origin of termites, the information on caste-biased gene and exon expression was extracted from the output produced in a previous study for *Z. nevadensis, C. secundus* and *M. natalensis* (Harrison et al., 2018). For *Reticulitermes speratus*, we used expression data from a previous study (Shigenobu et al., 2022) and ran differential gene expression analysis using the R package DESeq2 (Love et al., 2014). The proportions of caste-biased genes among non-rearranged and rearranged genes were compared using chisq.test function in R.

### CpGo/e and methylation analyses

To estimate the methylation level of DAs, the CpGo/e was calculated by comparing observed to expected CpG counts per domain where the expected values are the product of cytosine and guanine fractions. We calculated the weighted mean of CpGo/e per DA where the weights were the fractions of CG pairs. To test whether CpGo/e of domains rearranged in the origin of termites is lower than CpGo/e of non-rearranged domains, we performed a one-sided Wilcoxon test in R.

To test whether lower CpGo/e is associated with rearrangements, we extracted the ancestral arrangements that underwent changes in the origin of termites from the DomRates output. We excluded fusions, because some single domains that were fused in the origin of termites were over-represented, for example zinc finger domain (PF00096). Next, we calculated CpGo/e for all DAs as previously described and compared DAs that were changed in the origin of termites to the rest using one-sided Wilcoxon test in R.

Moreover, we used methylation data for *M. natalensis* from a previous study (Harrison et al., 2022) to test if methylation is indeed higher in rearranged genes. We computed average methylation values over gene regions (flank, cds, intron) and replicates for each gene and compared between genes rearranged in both origins of termites and true workers and non-rearranged genes using one-sided Wilcoxon test in R.

### Transposable elements in close proximity

Data on TE counts per gene was taken from a previous study for *Z. nevadensis, C. secundus, M. natalensis* and *B. germanica*, which included proportions of 10kb flanking regions (3’- and 5’-of genes) covered by TEs (Harrison et al., 2018). We compared the levels of TE content in flanking regions for each species between rearranged and non-rearranged genes. For *M. natalensis*, rearranged genes included genes rearranged at the origins of termites and true workers.

## Results

### Over 5000 domain rearrangements in Blattodea

Using proteomes of three non-social cockroaches, two termites with totipotent workers, three termites with true workers and two outgroup species (*Eriosoma lanigerum*, woolly apple aphid, and *Frankliniella occidentalis*, western flower thrips), we reconstructed protein domain rearrangement events along the Blattodea phylogeny. For this we annotated protein sequences with PFAM domains, version 30.0 (Mistry et al., 2021), and used DomRates (Dohmen et al., 2020) to infer domain architectures (DAs) of all ancestral nodes based on those of the analysed species, thus inferring all rearrangements at each node of the tree. Overall, we observed more than 5000 rearrangements in Blattodea, most of which were fusions of two ancestral DAs (44%), while the fission of one ancestral DA into two new DAs accounts for 13% of rearrangements. Moreover, we observed 22% and 18% of single domain and terminal domain losses, respectively. Novel domain emergences correspond to 3% of all events (Figure 1, Supplementary Table 2). As proteome completeness scores, based on the existence of conserved DAs measured with DOGMA (Dohmen et al., 2016), varied among the analysed species (Fig. S1), we did not consider domain losses in subsequent analyses as they could be due to incompleteness of the analysed genome assemblies or annotations.

**Figure 1:**
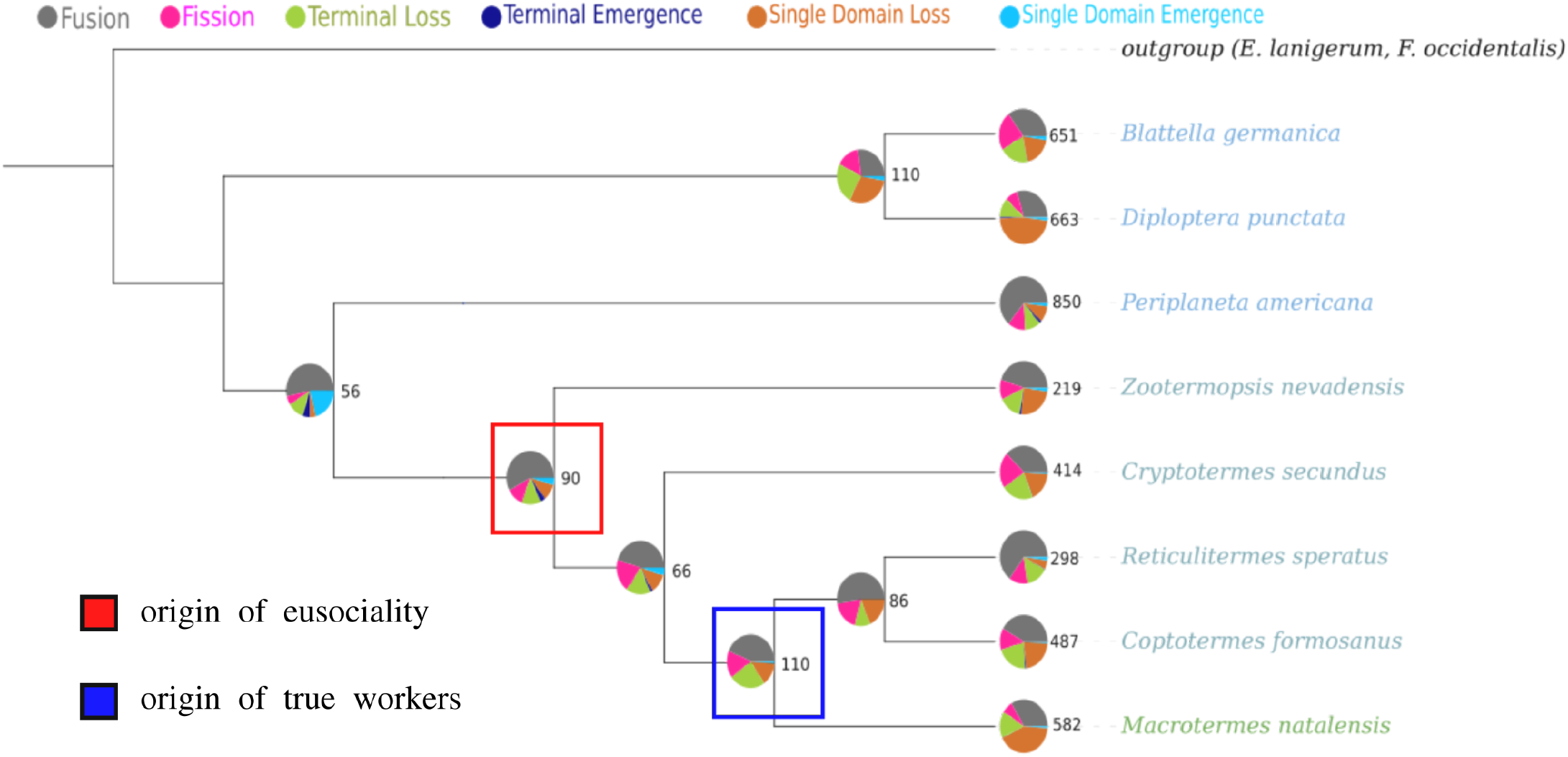
Domain rearrangements in Blattodea. For each node, the pie charts represent the fractions of different types of rearrangements shown above, and the numbers are the total amount of rearrangements. The nodes of the origin of termites and the origin of true workers are indicated with the red and blue squares, respectively.

### Domains related to longevity, nutrition and development rearranged at the root of termite eusociality

We were particularly interested in functional changes that occurred at two nodes: (i) the origin of termites where eusociality emerged (Figure 1, red box) and (ii) the origin of true workers associated with various novel traits such as foraging behaviour, differences in development, larger colonies and and extended lifespan of reproductive castes (Figure 1, blue box). At the origin of termite eusociality, we found 90 novel domain arrangements, including 52 fusions, 10 fissions and 4 and 3 single and terminal domain emergences, respectively. These seven ‘novel’ domains exist elsewhere in the tree of life but were inferred here as ‘emerged’ at the root of termites, since they were not found in any of the five analysed non-social species (three cockroaches, the aphid *E lanigerum* and the thrips *F. occidentalis*). These seven termite specific domains are associated with functions related to protein homeostasis (PF05025, PF01814 & PF07297), mitochondrial functions (PF10642), DNA repair (PF06331) and insulin production (PF11548) (Table 1). A further domain (PF11600) is found in the chromatin assembly factor 1 complex.

**Table 1:**
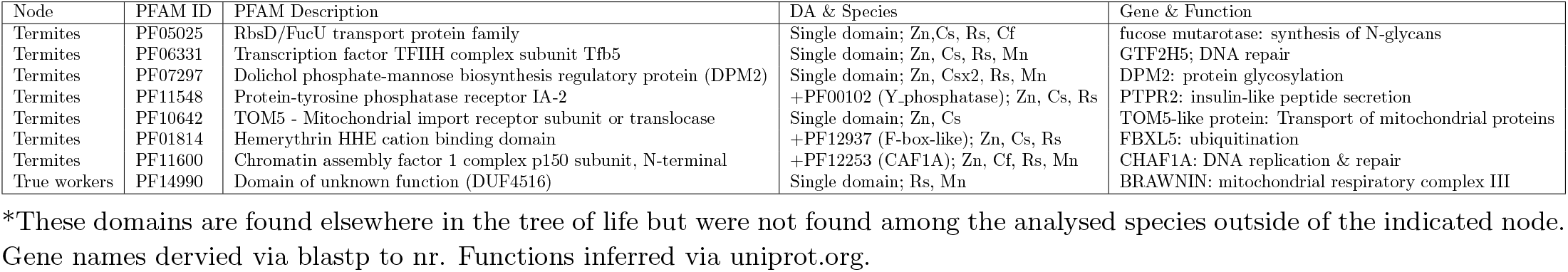
PFAM IDs and descriptions of domains classed as ‘emerged’ at the origin of termites and the origin of true workers in relation to other analysed species.

To understand the functions that were affected by domain rearrangements we performed two types of Gene Ontology (GO) term enrichment analyses. First, at all of the internal nodes of the phylogeny shown in Fig. 1, we compared the domains, and their associated GO-terms, involved in novel domain rearrangements to all DAs contained in the full species set. At the origin of termite eusociality (red box in Figure 1), we found an enrichment of various GO-terms related to development and metamorphosis, such as the regulation of embryonic development, cell migration and cell adhesion, as well as cilium assembly and movement (Fig. 2 A). Post-translational modifications of proteins also appear to be important for the evolution of termites, with protein phosphorylation and dolichol metabolism, involved in glycosylation (Carroll et al., 1992), both enriched at the root of termites. This is further supported by similar enriched GO-terms at other internal termite nodes but a lack thereof in cockroach nodes (Supplementary Figure 2). Finally, in each termite species, we analysed functional enrichment of genes containing DAs that were rearranged at the origin of termites. We found common GO-terms related to intracellular protein transport in three termites *Z. nevadensis, C. secundus*, and *R. speratus*. Several GO-terms involved in DNA repair mechanisms as well as stress response were found in *C. secundus, R. speratus*, and the higher termite *M. natalensis*.

**Figure 2:**
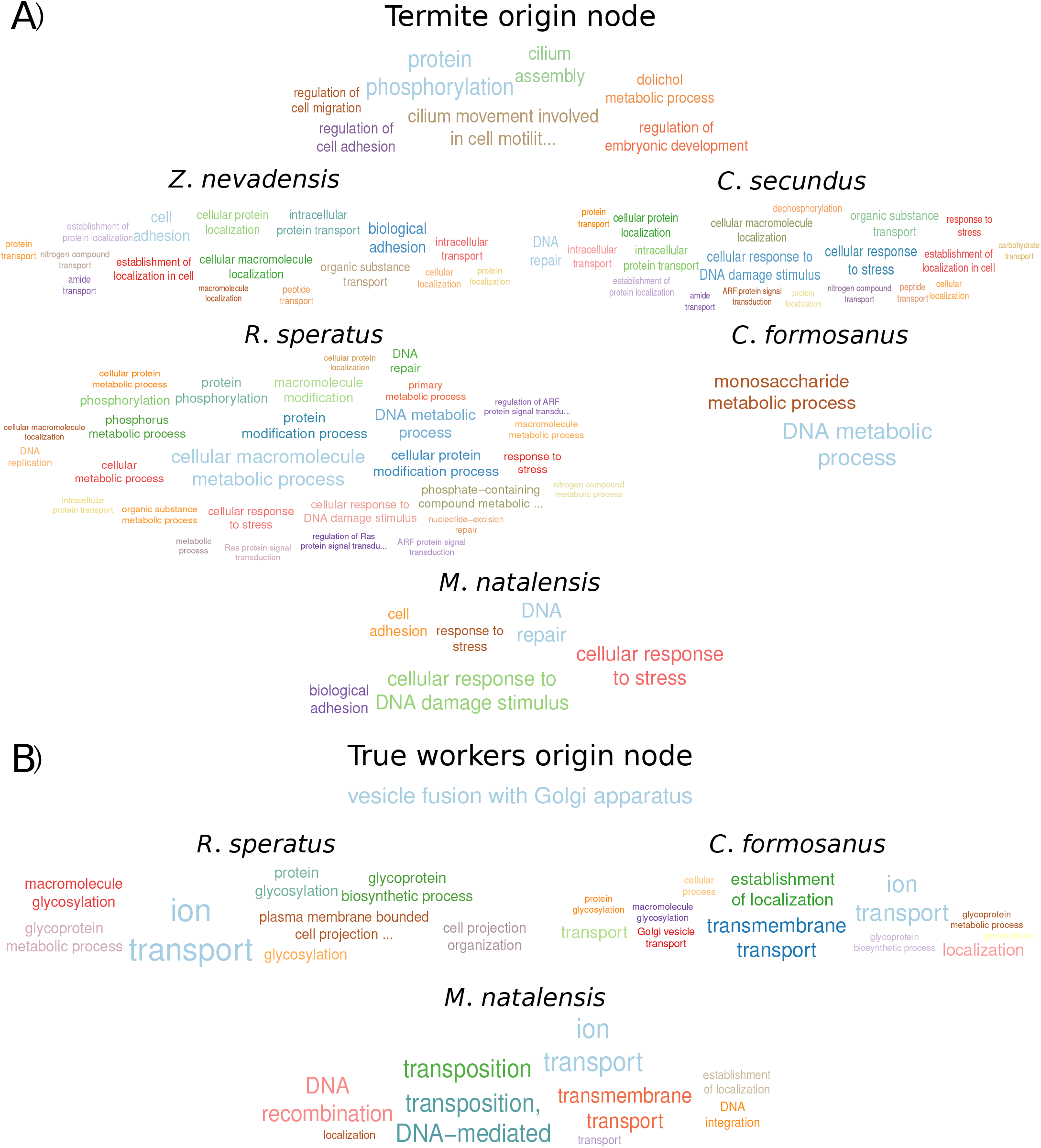
Enriched GO-terms for rearranged domains and affected genes for each termite species at two nodes. A: Enriched GO-terms at the ancestral node of termites, B: Enriched GO-terms at the node at which true workers emerged. Font sizes of GO-terms are relative to the inverse p-value. For full details see Table S3-11.

### Rearrangements enriched for protein glycosylation at the origin of true workers

At the origin of true workers (blue box in Fig. 1), we inferred 110 additional new DAs, including 48 fusions, 18 fissions and one single domain emergence. Within the ten analysed species, this ‘novel’ domain (PF14990) was found only in two of the three species with true workers, and is described as a domain of unknown function (DUF4516; Table 1). A protein containing this domain was found both in *M. natalensis* (Mnat 01442) and *R. speratus* (RS000595), both of which showed, via Blastp (Camacho et al., 2009), sequence similarity to the protein BRAWNIN with up to 68.75% and 66.67% identity.

The domains that were rearranged at the root of true workers (blue box in Fig. 1) were enriched for a single GO-term - “vesicle fusion with Golgi apparatus” (Fig. 2B). The related domain, PF04869 (Uso1 p115 head), and the corresponding protein, Golgi matrix protein p115, lost a terminal domain (PF04871, Uso1 p115 C) in *C. formosanus* and *M. natalensis* compared to lower termites and cockroaches. We analysed the proteins that were rearranged at the origin of workers in each species (Fig. 2B). The majority of enriched GO-terms were related to protein glycosylation, or protein and ion transport in the termites with true workers, while we also found an enrichment of transposition in *M. natalensis*.

### Genes with domain rearrangements show caste-biased gene and exon expression

We hypothesized that genes, in which domains were rearranged at the root of termite eusociality or at the emergence of true workers, were important for the evolution of castes. An increased importance of these rearranged genes for particular castes should therefore be reflected in caste-biased expression. To test this hypothesis, we compared proportions of caste-biased genes within rearranged and non-rearranged genes for the four species, for which caste-specific expression data were available. Confirming our expectations, we observed, with a *χ*-squared test, a significant enrichment of caste-biased genes within rearranged genes for *C. secundus* (44% of genes with rearrangements are caste-biased vs 26% of genes without rearrangements, p-value *<* 0.0001) and *M. natalensis* (75% of genes with rearrangements vs 39%, p-value *<* 0.00001). The proportion of caste-biased genes with domain rearrangements was also higher for *Z. nevadensis* (60% vs 56%, p-value = 0.519) and *R. flavipes* (11% vs 10%, p-value = 0.998), but the difference was not statistically significant.

Alternative splicing offers a further mechanism for gaining caste-specific protein functions (Harrison et al., 2022; Price et al., 2018). To test whether genes with domain rearrangements also have an increased rate of caste-biased alternative splicing, we used data on differential exon expression for *M. natalensis* from a previous study (Harrison et al., 2022). We observed an enrichment of genes alternatively spliced in any caste within the subset of rearranged genes compared to non-rearranged genes (Figure 3). The same pattern was observed separately for genes alternatively spliced in kings, alate queens and workers (Supplementary Figure 3a-c) but not in mature queens (Supplementary Figure 3d).

**Figure 3:**
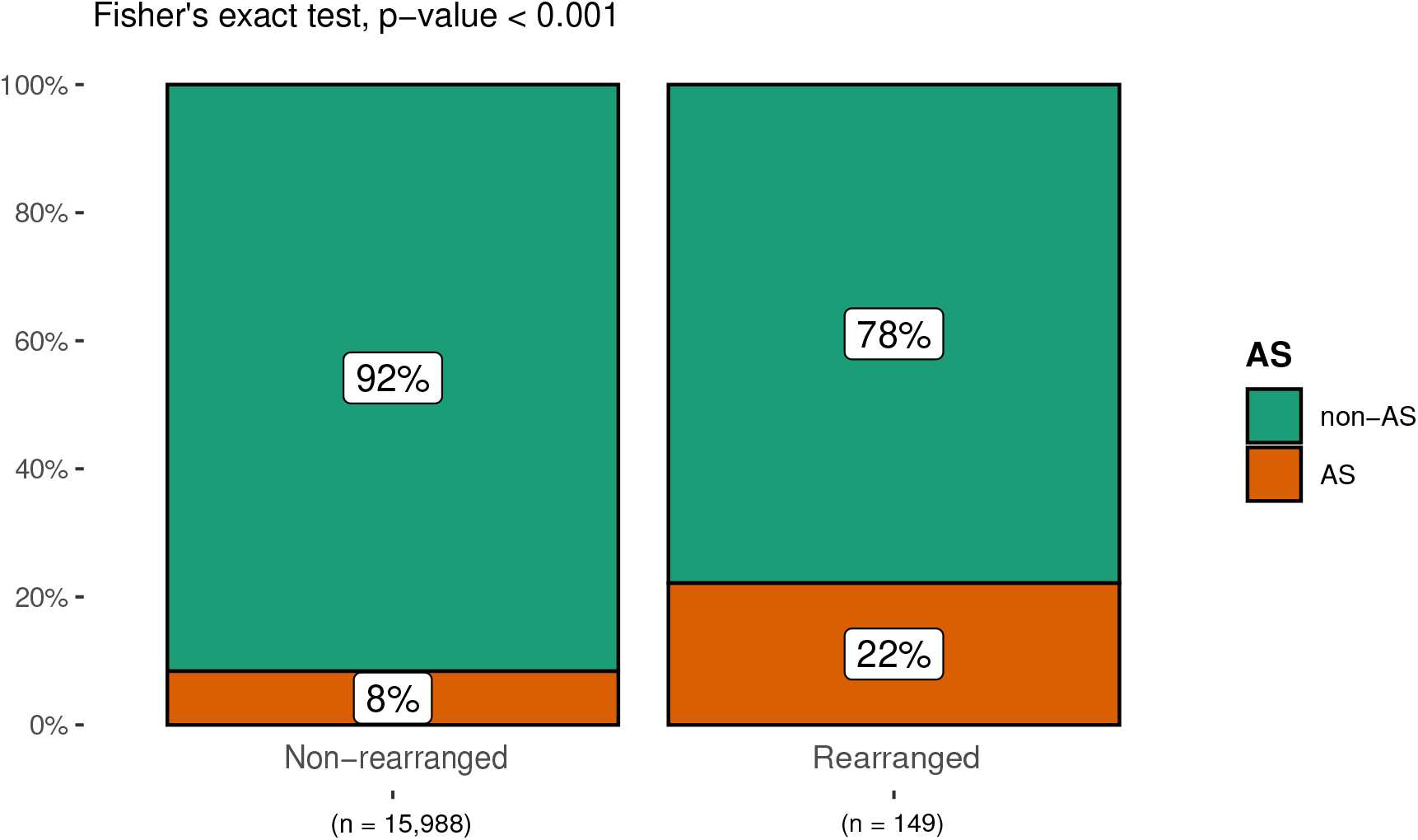
The proportions of alternatively spliced (AS) genes within rearranged and non-rearranged genes at the origins of termites and true workers in *M. natalensis*

### Genes with domain rearrangements have higher estimated methylation level

Because genes with domain rearrangements show different patterns of expression compared to non-rearranged genes, we expected to find differences in the regulation of their expression. To test this we compared levels of estimated methylation in genes with and without domain rearrangements in cockroaches and termites. For this we compared observed versus expected CpG counts, where a depletion is known to correlate with higher levels of DNA methylation (Park et al., 2011). Confirming expectations, we observed lower CpGo/e values for rearranged compared to non-rearranged genes in all analysed blattodean species, significantly in all except *R. speratus* (Table 2). This difference in CpGo/e indicates higher DNA methylation in rearranged genes.

To further test the causal link between the lower CpGo/e levels and domain rearrangements, we analysed genes with the ancestral DAs in cockroaches that underwent evolutionary changes in termites. In contrast to rearranged genes in termites, cockroach genes carrying the ancestral DAs did not significantly differ from other genes in terms of CpGo/e for all three species of cockroach (Supplementary table 3). To confirm the ability of CpGo/e to predict relative DNA methylation levels in rearranged genes, we analysed existing bisulfite sequencing data for *M. natalensis* (Harrison et al., 2022). Our estimations were confirmed. The rearranged genes showed significantly higher methylation levels compared to non-rearranged genes (p-value = 0.022, one-sided Wilcoxon test).

**Table 2:** CpGo/e in rearranged and non-rearranged domains. P-value for one-sided Wilcoxon test

| Species | N of RA | N of non-RA | mean CpGo/e<br>for RA | mean CpGo/e<br>for non-RA | p-value |
| --- | --- | --- | --- | --- | --- |
| *Z. nevadensis* | 105 | 9751 | 0.508 | 0.579 | 0.014 |
| *C. secundus* | 99 | 12311 | 0.448 | 0.535 | < 0.001 |
| *R. speratus* | 77 | 10439 | 0.512 | 0.538 | 0.250 |
| *C. formosanus* | 54 | 9779 | 0.446 | 0.536 | 0.011 |
| *M. natalensis* | 52 | 9295 | 0.440 | 0.568 | 0.002 |
| *B. germanica* | 301 | 11126 | 0.538 | 0.604 | < 0.001 |
| *D. punctata* | 386 | 13433 | 0.480 | 0.543 | < 0.001 |
| *P. americana* | 152 | 14693 | 0.553 | 0.656 | < 0.001 |

### Depletion of transposable elements in close proximity of rearranged genes in termites

To understand if domain rearrangements that occurred along the evolution of termites were due to TE activity, we analyzed TE counts in close proximity to the corresponding genes for the termites *Z. nevadensis, C. secundus* and *M. natalensis* (data from (Harrison et al., 2018), Supplementary Figure 4). We did not observe increased TE content in 10kb flanking regions of the genes rearranged at the origin of termites. In fact, TEs were depleted in rearranged genes compared to non-rearranged genes in all three termite species, significantly so in *Z. nevadensis* (median TE content: 0.23 vs. 0.27; p-value = 2.56×10^−3^) and *M. natalensis* for genes rearranged at the root of true workers (median TE content: 0.32 vs. 0.36; p-value = 6.91×10^−4^). In the cockroach *B. germanica*, on the other hand, TE content did not differ between rearranged and non-rearranged genes (median content: 0.46 vs. 0.46; Supplementary Figure 5).

## Discussion

Blattodea genomes show a reduction in proteome size and gene family contractions along with the evolution of eusociality in termites (Harrison et al., 2018). Here, we tested whether the functional novelty involved in the evolution of termites was acquired by changes in existing proteins. For this we analysed domain architectures (DAs) in the proteomes of all available Blattodea genomes (three cockroaches and five termites) and two outgroup species. We observed more than 5000 rearrangements in Blattodea with the proportions of different rearrangement events similar to previous studies (Dohmen et al., 2020; Thomas et al., 2020). At the origin of termites, when small wood-dwelling colonies emerged with totipotent workers and long-lived reproductives, we discovered 90 new domain arrangements that may be linked to the emergence of eusociality as well as a specialisation on wood feeding. Along with the transition to superorganismality, which involved the evolution of sterile, ‘true’ workers, larger, more complex, foraging colonies, and even longer-lived reproductives, we detected 110 further evolutionary changes in DAs. We uncovered seven novel domains at the origin of termites, i.e. not found in the analysed cockroaches and outgroups, but only one domain that, within our species set, was specific to termite species with true workers, indicating the transition to superorganismality mainly involved the rearrangement of existing DAs.

At the emergence of eusociality, novel domains affected proteins with functions that can be related to changes in energy consumption, diets and lifespan among castes, such as DNA replication and mitochondrial functions. These along with proteins with functions in protein homeostasis (glycosylation and ubiquitination), DNA repair and nutrient sensing (IIS) can be related to several important ageing mechanisms (López-Otín et al., 2013). These mechanisms have previously been linked to extreme longevity and high, maintained reproductive fitness in termite queens (Séité et al., 2022; Monroy Kuhn et al., 2019). Modifications in genes containing these domains may, therefore, have allowed the evolution of longevity in termite queens. A domain emergence in the chromatin assembly factor 1 complex at the root of termites may be important for remodelled developmental programmes necessary for the evolution of castes. Rearranged DAs at the origin of termites are enriched for several developmental GO-terms, such as regulation of embryonic development, cell adhesion and cell migration, which are likely linked to the evolution of novel caste phenotypes in the termites. As in the novel domains, we also found DA rearrangments in termites related to DNA repair and response to stress which may be related to the high longevity in termite queens.

The domain that was unique to termites with true workers in our data set has a putative role in the mitochondrial respiratory chain within the gene BRAWNIN, which may be linked to increased caste-specific differences in energy consumption due to foraging of workers and increased longevity and reproductive output of kings and queens. A single GO-term was enriched at the node where in our species set true workers emerged. The domain (F04869: Uso1 p115 head) that was associated with this enriched GO-term, “vesicle fusion with Golgi apparatus", is found in the gene p115, which is important for correct morphology and organisation of the Golgi apparatus and transitional endoplasmic reticulum (Kondylis and Rabouille, 2003). Further domain rearrangements at the origin of true workers, indicated the importance of protein glycosylation and protein transport in larger, more socially complex termite colonies.

One of the characteristics of eusociality is the division of labour with genomes producing distinct castes that differ in morphology, behaviour and physiology. These differences can be reflected in caste-specific patterns of gene expression and splicing (Scharf et al., 2005; Terrapon et al., 2014; Harrison et al., 2018, 2022). We observed an enrichment of differentially expressed and spliced genes amongst genes with domain rearrangements indicating the potential role of rearranged genes in the evolution of castes. One of the mechanisms shaping this pattern might be methylation as we observed an increase in gene body methylation for rearranged compared to non-rearranged genes using bisulfite sequencing data for *M. natalensis* and an estimate of methylation: CpGo/e. That we find no differences in CpGo/e between gene sets containing the ancestral DAs of these rearranged genes, indicates changes in methylation may have occurred after rearrangements took place. The observed differences in methylation for rearranged and non-rearranged genes indicate potential changes in the regulation of these sets of genes. Overall, the results on differential patterns of expression and splicing of rearranged genes in termites suggest the importance of domain rearrangements in the evolution of termite castes and the potential properties of genes involved in the evolution of eusociality in termites. However, increased methylation might be a common feature of rearranged genes as we found this pattern in both termites and cockroaches.

Some genomic innovations, including domain rearrangements, are associated with the activity of transposable elements (TEs) (Bourque et al., 2018; Buljan et al., 2010). However, we observed a depletion of TEs in the close proximity of the genes with domain rearrangements that occurred in the origins of termites and true workers in *Z. nevadensis* and *M. natalensis*. This might indicate other potential mechanisms involved in the domain rearrangements in termites such as recombination, as well as possible selection against TE insertions as an indicator of the functional importance of these genes (Bartolomé et al., 2002; Kent et al., 2017).

Altogether, our findings support a role of domain rearrangements in the evolution of termite castes with rearranged genes playing an important role in caste-biased gene expression and splicing. Domain rearrangements affected functions related to queen longevity and major changes in development. Our analyses indicate the greatest functional novelty due to domain evolution already arose at the origin of eusociality, where simple societies emerged with totipotent workers. The emergence of true workers along with increasing social complexity, such as larger foraging colonies and even longer-lived reproductives, on the other hand, was accompanied by more domain rearrangements, within similar functional categories. These analyses incorporated all currently available blattodean genomes. Further genomes will allow us to better pinpoint domain novelties and relate these to the emergence and elaboration of specific social traits such as worker sterility, foraging, queen lifespan and colony size, and differentiate these from confounding ecological traits, such as wood-feeding.

Important lineages in this respect are the subsocial sister group of all termites, the wood-dwelling *Cryptocercus* or the rather ancient lineage of *Mastotermes*, in which large foraging colonies with true workers evolved independently to the termites studied in the present study. Nevertheless, the observations reported here already suggest an important role for domain rearrangements in the evolution of termite eusociality with intriguing implications for other origins of eusociality, in which this source of protein novelty is so far under appreciated.

## Acknowledgements

AM is supported by the DFG grant HA 8997/1-1 to MH. The authors declare no conflicts of interest.

## Author contributions

MH conceived the study. AM carried out all analyses and wrote the first manuscript draft. ED & MH assisted with analyses and interpreting results. All authors worked on revising the manuscript.

## Data accessibility

All analyses were carried out on publicly available data, using published tools that are cited within the manuscript. Inquiries and requests can be directed to the corresponding author.

## References

Abe, T. (1990). Evolution of worker caste in termites. Social insects and the environments, pages 29–30.

Alexa, A., Rahnenfuhrer, J., et al. (2010). topgo: enrichment analysis for gene ontology. R package version 2.38.1, 2(0):2010.

Bartolomé, C., Maside, X., and Charlesworth, B. (2002). On the abundance and distribution of transposable elements in the genome of drosophila melanogaster. Molecular biology and evolution, 19(6):926–937.

Biello, R., Singh, A., Godfrey, C. J., Fernández, F. F., Mugford, S. T., Powell, G., Hogenhout, S. A., and Mathers, T. C. (2021). A chromosome-level genome assembly of the woolly apple aphid, eriosoma lanigerum hausmann (hemiptera: Aphididae). Molecular ecology resources, 21(1):316–326.

Bourque, G., Burns, K. H., Gehring, M., Gorbunova, V., Seluanov, A., Hammell, M., Imbeault, M., Izsvák, Z., Levin, H. L., Macfarlan, T. S., et al. (2018). Ten things you should know about transposable elements. Genome biology, 19:1–12.

Buljan, M., Frankish, A., and Bateman, A. (2010). Quantifying the mechanisms of domain gain in animal proteins. Genome biology, 11(7):1–15.

Camacho, C., Coulouris, G., Avagyan, V., Ma, N., Papadopoulos, J., Bealer, K., and Madden, T. L. (2009). Blast+: architecture and applications. BMC bioinformatics, 10:1–9.

Carroll, K., Guthrie, N., and Ravi, K. (1992). Dolichol: function, metabolism, and accumulation in human tissues. Biochemistry and Cell Biology, 70(6):382–384.

Coban, A., Bornberg-Bauer, E., and Kemena, C. (2022). Domain evolution of vertebrate blood coagulation cascade proteins. Journal of Molecular Evolution, 90(6):418–428.

Crespi, B. J. (1996). Comparative analysis of the origins and losses of eusociality: causal mosaics and historical uniqueness. Phylogenies and the comparative method in animal behavior, pages 253–287.

Dohmen, E., Klasberg, S., Bornberg-Bauer, E., Perrey, S., and Kemena, C. (2020). The modular nature of protein evolution: domain rearrangement rates across eukaryotic life. BMC Evolutionary Biology, 20(1):30.

Dohmen, E., Kremer, L. P., Bornberg-Bauer, E., and Kemena, C. (2016). Dogma: domain-based transcriptome and proteome quality assessment. Bioinformatics, 32(17):2577–2581.

Ekman, D., Björklund, Å. K., and Elofsson, A. (2007). Quantification of the elevated rate of domain rearrangements in metazoa. Journal of molecular biology, 372(5):1337–1348.

Elsner, D., Meusemann, K., and Korb, J. (2018). Longevity and transposon defense, the case of termite reproductives. Proceedings of the National Academy of Sciences, 115(21):5504–5509.

Emms, D. M. and Kelly, S. (2019). Orthofinder: phylogenetic orthology inference for comparative genomics. Genome biology, 20(1):1–14.

Finn, R. D., Coggill, P., Eberhardt, R. Y., Eddy, S. R., Mistry, J., Mitchell, A. L., Potter, S. C., Punta, M., Qureshi, M., Sangrador-Vegas, A., et al. (2016). The pfam protein families database: towards a more sustainable future. Nucleic acids research, 44(D1):D279–D285.

Forslund, K. and Sonnhammer, E. L. (2012). Evolution of protein domain architectures. Evolutionary Genomics: Statistical and Computational Methods, Volume 2, pages 187–216.

Forslund, S. K., Kaduk, M., and Sonnhammer, E. L. (2019). Evolution of protein domain architectures. Evolutionary genomics: statistical and computational methods, pages 469–504.

Fouks, B., Harrison, M. C., Mikhailova, A. A., Marchal, E., English, S., Carruthers, M., Jennings, E. C., Pippel, M., Attardo, G. M., Benoit, J. B., et al. (2022). Live-bearing cockroach genome reveals convergent evolutionary mechanisms linked to viviparity in insects and beyond. bioRxiv, pages 2022–02.

Harrison, M. C., Dohmen, E., George, S., Sillam-Dussès, D., Séité, S., and Vasseur-Cognet, M. (2022). Complex regulatory role of dna methylation in caste-and age-specific expression of a termite. Open Biology, 12(7):220047.

Harrison, M. C., Jongepier, E., Robertson, H. M., Arning, N., Bitard-Feildel, T., Chao, H., Childers, C. P., Dinh, H., Doddapaneni, H., Dugan, S., et al. (2018). Hemimetabolous genomes reveal molecular basis of termite eusociality. Nature ecology & evolution, 2(3):557–566.

Inward, D., Beccaloni, G., and Eggleton, P. (2007). Death of an order: a comprehensive molecular phylogenetic study confirms that termites are eusocial cockroaches. Biology letters, 3(3):331–335.

Itakura, S., Yoshikawa, Y., Togami, Y., and Umezawa, K. (2020). Draft genome sequence of the termite, coptotermes formosanus: Genetic insights into the pyruvate dehydrogenase complex of the termite. Journal of Asia-Pacific Entomology, 23(3):666–674.

Itoh, M., Nacher, J. C., Kuma, K.-i., Goto, S., and Kanehisa, M. (2007). Evolutionary history and functional implications of protein domains and their combinations in eukaryotes. Genome biology, 8:1–15.

Kaib, M., Hacker, M., and Brandl, R. (2001). Egg-laying in monogynous and polygynous colonies of the termite macrotermes michaelseni (isoptera, macrotermitidae). Insectes sociaux, 48:231–237.

Kapheim, K. M., Pan, H., Li, C., Salzberg, S. L., Puiu, D., Magoc, T., Robertson, H. M., Hudson, M. E., Venkat, A., Fischman, B. J., et al. (2015). Genomic signatures of evolutionary transitions from solitary to group living. Science, 348(6239):1139–1143.

Keller, L. (1998). Queen lifespan and colony characteristics in ants and termites. Insectes sociaux, 45:235–246.

Kemena, C., Dohmen, E., and Bornberg-Bauer, E. (2019). Dogma: a web server for proteome and transcriptome quality assessment. Nucleic Acids Research, 47(W1):W507–W510.

Kent, D. and Simpson, J. (1992). Eusociality in the beetle austroplatypus incompertus (coleoptera: Curculionidae). Naturwissenschaften, 79(2):86–87.

Kent, T. V., Uzunović, J., and Wright, S. I. (2017). Coevolution between transposable elements and recombination. Philosophical Transactions of the Royal Society B: Biological Sciences, 372(1736):20160458.

Kondylis, V. and Rabouille, C. (2003). A novel role for dp115 in the organization of ter sites in drosophila. The Journal of cell biology, 162(2):185–198.

Korb, J. and Hartfelder, K. (2008). Life history and development-a framework for understanding developmental plasticity in lower termites. Biological Reviews, 83(3):295–313.

Korb, J. and Lenz, M. (2004). Reproductive decision-making in the termite, cryptotermes secundus (kalotermitidae), under variable food conditions. Behavioral Ecology, 15(3):390–395.

Korb, J. and Thorne, B. (2017). Sociality in termites. Comparative social evolution, pages 124–153.

Legendre, F., Whiting, M. F., Bordereau, C., Cancello, E. M., Evans, T. A., and Grandcolas, P. (2008). The phylogeny of termites (dictyoptera: Isoptera) based on mitochondrial and nuclear markers: implications for the evolution of the worker and pseudergate castes, and foraging behaviors. Molecular phylogenetics and evolution, 48(2):615–627.

Li, S., Zhu, S., Jia, Q., Yuan, D., Ren, C., Li, K., Liu, S., Cui, Y., Zhao, H., Cao, Y., et al. (2018). The genomic and functional landscapes of developmental plasticity in the american cockroach. Nature communications, 9(1):1008.

Lo, N., Engel, M. S., Cameron, S., Nalepa, C. A., Tokuda, G., Grimaldi, D., Kitade, O., Krishna, K., Klass, K.-D., Maekawa, K., et al. (2007). Save isoptera: a comment on inward et al. Biology letters, 3(5):562–563.

López-Otín, C., Blasco, M. A., Partridge, L., Serrano, M., and Kroemer, G. (2013). The hallmarks of aging. Cell, 153(6):1194–1217.

Love, M. I., Huber, W., and Anders, S. (2014). Moderated estimation of fold change and dispersion for rna-seq data with deseq2. Genome biology, 15(12):1–21.

Meyer, V., Crewe, R., Braack, L., Groeneveld, H., and Van der Linde, M. (2000). Intracolonial demography of the moundbuilding termite macrotermes natalensis (haviland)(isoptera, termitidae) in the northern kruger national park, south africa. Insectes Sociaux, 47:390–397.

Mistry, J., Chuguransky, S., Williams, L., Qureshi, M., Salazar, G. A., Sonnhammer, E. L., Tosatto, S. C., Paladin, L., Raj, S., Richardson, L. J., et al. (2021). Pfam: The protein families database in 2021. Nucleic acids research, 49(D1):D412–D419.

Mitchell, A., Chang, H.-Y., Daugherty, L., Fraser, M., Hunter, S., Lopez, R., McAnulla, C., McMenamin, C., Nuka, G., Pesseat, S., et al. (2015). The interpro protein families database: the classification resource after 15 years. Nucleic acids research, 43(D1):D213–D221.

Monroy Kuhn, J. M., Meusemann, K., and Korb, J. (2019). Long live the queen, the king and the commoner? transcript expression differences between old and young in the termite cryptotermes secundus. PLoS One, 14(2):e0210371.

Moore, A. D., Björklund, Å. K., Ekman, D., Bornberg-Bauer, E., and Elofsson, A. (2008). Arrangements in the modular evolution of proteins. Trends in biochemical sciences, 33(9):444–451.

Noirot, C. (1988). The worker caste is polyphyletic in termites. Sociobiology, 14:15–20.

Noirot, C. and Pasteels, J. (1987). Ontogenetic development and evolution of the worker caste in termites. Experientia, 43:851–860.

Park, J., Peng, Z., Zeng, J., Elango, N., Park, T., Wheeler, D., Werren, J. H., and Yi, S. V. (2011). Comparative analyses of dna methylation and sequence evolution using nasonia genomes. Molecular biology and evolution, 28(12):3345–3354.

Patthy, L. (1985). Evolution of the proteases of blood coagulation and fibrinolysis by assembly from modules. Cell, 41(3):657–663.

Patthy, L. (2003). Modular assembly of genes and the evolution of new functions. Genetica, 118:217–231.

Pawson, T. (1995). Protein modules and signalling networks. Nature, 373(6515):573–580.

Poulsen, M., Hu, H., Li, C., Chen, Z., Xu, L., Otani, S., Nygaard, S., Nobre, T., Klaubauf, S., Schindler, P. M., et al. (2014). Complementary symbiont contributions to plant decomposition in a fungus-farming termite. Proceedings of the National Academy of Sciences, 111(40):14500–14505.

Price, J., Harrison, M., Hammond, R., Adams, S., Gutierrez-Marcos, J., and Mallon, E. (2018). Alternative splicing associated with phenotypic plasticity in the bumble bee bombus terrestris. Molecular Ecology, 27(4):1036–1043.

Pull, C. D. and McMahon, D. P. (2020). Superorganism immunity: a major transition in immune system evolution. Frontiers in Ecology and Evolution, page 186.

Rehan, S. M. and Toth, A. L. (2015). Climbing the social ladder: the molecular evolution of sociality. Trends in Ecology & Evolution, 30(7):426–433.

Roisin, Y. and Korb, J. (2011). Social organisation and the status of workers in termites. Biology of termites: a modern synthesis, pages 133–164.

Rotenberg, D., Baumann, A. A., Ben-Mahmoud, S., Christiaens, O., Dermauw, W., Ioannidis, P., Jacobs, C. G., Vargas Jentzsch, I. M., Oliver, J. E., Poelchau, M. F., et al. (2020). Genome-enabled insights into the biology of thrips as crop pests. BMC biology, 18(1):1–37.

Scharf, M., Wu-Scharf, D., Zhou, X., Pittendrigh, B., and Bennett, G. (2005). Gene expression profiles among immature and adult reproductive castes of the termite reticulitermes flavipes. Insect molecular biology, 14(1):31–44.

Séité, S., Harrison, M. C., Sillam-Dussès, D., Lupoli, R., Van Dooren, T. J., Robert, A., Poissonnier, L.-A., Lemainque, A., Renault, D., Acket, S., et al. (2022). Lifespan prolonging mechanisms and insulin upregulation without fat accumulation in long-lived reproductives of a higher termite. Communications Biology, 5(1):44.

Shell, W. A., Steffen, M. A., Pare, H. K., Seetharam, A. S., Severin, A. J., Toth, A. L., and Rehan, S. M. (2021). Sociality sculpts similar patterns of molecular evolution in two independently evolved lineages of eusocial bees. Communications Biology, 4(1):1–9.

Sherman, P. W., Jarvis, J. U., and Alexander, R. D. (2017). The biology of the naked mole-rat, volume 54. Princeton University Press.

Shigenobu, S., Hayashi, Y., Watanabe, D., Tokuda, G., Hojo, M. Y., Toga, K., Saiki, R., Yaguchi, H., Masuoka, Y., Suzuki, R., et al. (2022). Genomic and transcriptomic analyses of the subterranean termite reticulitermes speratus: Gene duplication facilitates social evolution. Proceedings of the National Academy of Sciences, 119(3):e2110361119.

Simola, D. F., Wissler, L., Donahue, G., Waterhouse, R. M., Helmkampf, M., Roux, J., Nygaard, S., Glastad, K. M., Hagen, D. E., Viljakainen, L., et al. (2013). Social insect genomes exhibit dramatic evolution in gene composition and regulation while preserving regulatory features linked to sociality. Genome research, 23(8):1235–1247.

Szathmáry, E. and Smith, J. M. (1995). The major transitions in evolution. WH Freeman Spektrum Oxford, UK:.

Terrapon, N., Li, C., Robertson, H. M., Ji, L., Meng, X., Booth, W., Chen, Z., Childers, C. P., Glastad, K. M., Gokhale, K., et al. (2014). Molecular traces of alternative social organization in a termite genome. Nature communications, 5(1):3636.

Thomas, G. W., Dohmen, E., Hughes, D. S., Murali, S. C., Poelchau, M., Glastad, K., Anstead, C. A., Ayoub, N. A., Batterham, P., Bellair, M., et al. (2020). Gene content evolution in the arthropods. Genome biology, 21(1):1–14.

Thorne, B. L., Breisch, N. L., and Haverty, M. I. (2002). Longevity of kings and queens and first time of production of fertile progeny in dampwood termite (isoptera; termopsidae; zootermopsis) colonies with different reproductive structures. Journal of Animal Ecology, 71(6):1030–1041.

Wilson, E. O. and Hölldobler, B. (2005). Eusociality: origin and consequences. Proceedings of the National Academy of Sciences, 102(38):13367–13371.

