## Supplementary Materials for "Major changes in domain arrangements are associated with the evolution of termite castes"

Alina A. Mikhailova[a],
Elias Dohmen[a],
Mark C. Harrison[a*]

[a] Institute for Evolution and Biodiversity, University of Münster
[*] Correspondence to m.harrison[at]uni-muenster.de

May 15, 2023

| Species | Completeness score, % |
|---|---|
| *E. lanigerum* | 75.72 |
| *F. occidentalis* | 86.16 |
| *B. germanica* | 59.70 |
| *D. punctata* | 68.82 |
| *P. americana* | 84.61 |
| *Z. nevadensis* | 90.01 |
| *C. secundus* | 83.64 |
| *R. speratus* | 93.88 |
| *C. formosanus* | 84.97 |
| *M. natalensis* | 75.66 |

Table 1: Proteome completeness score measured with DOGMA

| Node ID | Fusions | Fissions | TerminalLosses | TerminalEmergences | SingleDomainLosses | SingleDomainEmergences |
|---|---|---|---|---|---|---|
| 3 | 30 | 17 | 27 | 0 | 33 | 3 |
| 4 | 230 | 153 | 125 | 1 | 126 | 16 |
| 5 | 198 | 56 | 77 | 5 | 314 | 13 |
| 6 | 30 | 3 | 6 | 3 | 2 | 12 |
| 7 | 52 | 10 | 12 | 3 | 9 | 4 |
| 8 | 99 | 26 | 32 | 3 | 54 | 5 |
| 9 | 30 | 13 | 11 | 1 | 8 | 3 |
| 10 | 157 | 89 | 89 | 1 | 74 | 4 |
| 11 | 48 | 18 | 27 | 0 | 16 | 1 |
| 12 | 196 | 42 | 93 | 1 | 244 | 6 |
| 13 | 45 | 16 | 9 | 0 | 16 | 0 |
| 14 | 204 | 64 | 102 | 4 | 110 | 3 |
| 15 | 193 | 38 | 44 | 1 | 16 | 6 |
| 16 | 543 | 103 | 89 | 14 | 84 | 17 |

Table 2: Number of the reconstructed domain rearrangement types per each node (node numbers from Supplementary Figure 1)

| Node | GO.ID | Term | Annotated | Significant | Expected | fisher |
|---|---|---|---|---|---|---|
| 3 | GO:0045892 | negative regulation of transcription, DN... | 2 | 2 | 0.05 | 0.00066 |
| 3 | GO:0006412 | translation | 81 | 8 | 2.1 | 0.00251 |
| 3 | GO:0006298 | mismatch repair | 25 | 3 | 0.65 | 0.02529 |
| 3 | GO:0000387 | spliceosomal snRNP assembly | 1 | 1 | 0.03 | 0.02597 |
| 6 | GO:0006526 | arginine biosynthetic process | 1 | 1 | 0.01 | 0.009 |
| 6 | GO:0006555 | methionine metabolic process | 1 | 1 | 0.01 | 0.009 |
| 6 | GO:0006139 | nucleobase-containing compound metabolic... | 479 | 6 | 4.33 | 0.016 |
| 6 | GO:0006384 | transcription initiation from RNA polyme... | 2 | 1 | 0.02 | 0.018 |
| 6 | GO:0000723 | telomere maintenance | 5 | 1 | 0.05 | 0.044 |
| 7 | GO:0006468 | protein phosphorylation | 119 | 6 | 1.75 | 0.0061 |
| 7 | GO:0060271 | cilium assembly | 1 | 1 | 0.01 | 0.0147 |
| 7 | GO:0060294 | cilium movement involved in cell motilit... | 1 | 1 | 0.01 | 0.0147 |
| 7 | GO:0045995 | regulation of embryonic development | 2 | 1 | 0.03 | 0.0292 |
| 7 | GO:0019348 | dolichol metabolic process | 2 | 1 | 0.03 | 0.0292 |
| 7 | GO:0030155 | regulation of cell adhesion | 2 | 1 | 0.03 | 0.0292 |
| 7 | GO:0030334 | regulation of cell migration | 3 | 1 | 0.04 | 0.0434 |
| 9 | GO:0006260 | DNA replication | 29 | 3 | 0.46 | 0.0098 |
| 9 | GO:0000256 | allantoin catabolic process | 1 | 1 | 0.02 | 0.0158 |
| 9 | GO:0045454 | cell redox homeostasis | 2 | 1 | 0.03 | 0.0314 |
| 9 | GO:0010835 | regulation of protein ADP-ribosylation | 3 | 1 | 0.05 | 0.0467 |
| 9 | GO:0006360 | transcription by RNA polymerase I | 3 | 1 | 0.05 | 0.0467 |
| 11 | GO:0048280 | vesicle fusion with Golgi apparatus | 1 | 1 | 0.03 | 0.026 |
| 13 | GO:0006487 | protein N-linked glycosylation | 6 | 2 | 0.11 | 0.0045 |
| 13 | GO:0015986 | proton motive force-driven ATP synthesis | 10 | 2 | 0.18 | 0.0130 |
| 13 | GO:0043486 | histone exchange | 1 | 1 | 0.02 | 0.0181 |
| 13 | GO:0035556 | intracellular signal transduction | 68 | 4 | 1.23 | 0.0283 |
| 13 | GO:0006744 | ubiquinone biosynthetic process | 2 | 1 | 0.04 | 0.0358 |
| 13 | GO:0007018 | microtubule-based movement | 42 | 3 | 0.76 | 0.0382 |
| 13 | GO:0007165 | signal transduction | 175 | 9 | 3.16 | 0.0462 |

Table 3: GO enrichment analysis for novel domain arrangements per each inner node (node numbers from Supplementary Figure 1)

| GO.ID | Term | Annotated | Significant | Expected | pvalue |
|---|---|---|---|---|---|
| GO:0007155 | cell adhesion | 48 | 4 | 0.56 | 0.0022 |
| GO:0022610 | biological adhesion | 48 | 4 | 0.56 | 0.0022 |
| GO:0006886 | intracellular protein transport | 65 | 4 | 0.76 | 0.0067 |
| GO:0034613 | cellular protein localization | 68 | 4 | 0.8 | 0.0079 |
| GO:0070727 | cellular macromolecule localization | 68 | 4 | 0.8 | 0.0079 |
| GO:0071702 | organic substance transport | 107 | 5 | 1.25 | 0.0079 |
| GO:0046907 | intracellular transport | 73 | 4 | 0.85 | 0.0101 |
| GO:0051649 | establishment of localization in cell | 74 | 4 | 0.87 | 0.0106 |
| GO:0051641 | cellular localization | 86 | 4 | 1.01 | 0.0177 |
| GO:0015031 | protein transport | 87 | 4 | 1.02 | 0.0183 |
| GO:0015833 | peptide transport | 89 | 4 | 1.04 | 0.0198 |
| GO:0045184 | establishment of protein localization | 89 | 4 | 1.04 | 0.0198 |
| GO:0042886 | amide transport | 90 | 4 | 1.05 | 0.0205 |
| GO:0071705 | nitrogen compound transport | 93 | 4 | 1.09 | 0.0229 |
| GO:0008104 | protein localization | 94 | 4 | 1.1 | 0.0237 |
| GO:0033036 | macromolecule localization | 102 | 4 | 1.19 | 0.0308 |

Table 4: GO enrichment analysis for genes with domain rearrangements at the origin of termites in *Z. nevadensis*

| GO.ID | Term | Annotated | Significant | Expected | pvalue |
|---|---|---|---|---|---|
| GO:0006281 | DNA repair | 47 | 3 | 0.4 | 0.0071 |
| GO:0006974 | cellular response to DNA damage stimulus | 48 | 3 | 0.4 | 0.0075 |
| GO:0033554 | cellular response to stress | 48 | 3 | 0.4 | 0.0075 |
| GO:0071702 | organic substance transport | 105 | 4 | 0.88 | 0.0115 |
| GO:0006886 | intracellular protein transport | 60 | 3 | 0.51 | 0.0138 |
| GO:0034613 | cellular protein localization | 64 | 3 | 0.54 | 0.0164 |
| GO:0070727 | cellular macromolecule localization | 64 | 3 | 0.54 | 0.0164 |
| GO:0046907 | intracellular transport | 67 | 3 | 0.56 | 0.0185 |
| GO:0051649 | establishment of localization in cell | 68 | 3 | 0.57 | 0.0193 |
| GO:0006950 | response to stress | 74 | 3 | 0.62 | 0.0241 |
| GO:0016311 | dephosphorylation | 31 | 2 | 0.26 | 0.0277 |
| GO:0051641 | cellular localization | 81 | 3 | 0.68 | 0.0304 |
| GO:0015031 | protein transport | 83 | 3 | 0.7 | 0.0323 |
| GO:0015833 | peptide transport | 85 | 3 | 0.72 | 0.0344 |
| GO:0045184 | establishment of protein localization | 85 | 3 | 0.72 | 0.0344 |
| GO:0042886 | amide transport | 87 | 3 | 0.73 | 0.0365 |
| GO:0071705 | nitrogen compound transport | 88 | 3 | 0.74 | 0.0375 |
| GO:0008104 | protein localization | 91 | 3 | 0.77 | 0.0408 |
| GO:0008643 | carbohydrate transport | 5 | 1 | 0.04 | 0.0414 |
| GO:0032011 | ARF protein signal transduction | 5 | 1 | 0.04 | 0.0414 |
| GO:0032012 | regulation of ARF protein signal transdu... | 5 | 1 | 0.04 | 0.0414 |

Table 5: GO enrichment analysis for genes with domain rearrangements at the origin of termites in *C. secundus*

| GO.ID | Term | Annotated | Significant | Expected | pvalue |
|---|---|---|---|---|---|
| GO:0044260 | cellular macromolecule metabolic process | 770 | 13 | 4.53 | 3.5e-05 |
| GO:0006259 | DNA metabolic process | 118 | 5 | 0.69 | 0.00045 |
| GO:0006464 | cellular protein modification process | 406 | 8 | 2.39 | 0.00116 |
| GO:0036211 | protein modification process | 406 | 8 | 2.39 | 0.00116 |
| GO:0006468 | protein phosphorylation | 234 | 6 | 1.38 | 0.00159 |
| GO:0043412 | macromolecule modification | 434 | 8 | 2.55 | 0.00182 |
| GO:0016310 | phosphorylation | 242 | 6 | 1.42 | 0.00189 |
| GO:0006281 | DNA repair | 52 | 3 | 0.31 | 0.00315 |
| GO:0006793 | phosphorus metabolic process | 371 | 7 | 2.18 | 0.00350 |
| GO:0006796 | phosphate-containing compound metabolic ... | 371 | 7 | 2.18 | 0.00350 |
| GO:0006974 | cellular response to DNA damage stimulus | 59 | 3 | 0.35 | 0.00452 |
| GO:0033554 | cellular response to stress | 61 | 3 | 0.36 | 0.00497 |
| GO:0044237 | cellular metabolic process | 1405 | 14 | 8.26 | 0.00599 |
| GO:0006950 | response to stress | 69 | 3 | 0.41 | 0.00702 |
| GO:0044238 | primary metabolic process | 1685 | 15 | 9.91 | 0.01191 |
| GO:0006260 | DNA replication | 33 | 2 | 0.19 | 0.01548 |
| GO:0043170 | macromolecule metabolic process | 1384 | 13 | 8.14 | 0.01885 |
| GO:0071704 | organic substance metabolic process | 1754 | 15 | 10.31 | 0.01911 |
| GO:0044267 | cellular protein metabolic process | 635 | 8 | 3.73 | 0.01985 |
| GO:0006289 | nucleotide-excision repair | 5 | 1 | 0.03 | 0.02908 |
| GO:0007265 | Ras protein signal transduction | 6 | 1 | 0.04 | 0.03480 |
| GO:0032011 | ARF protein signal transduction | 6 | 1 | 0.04 | 0.03480 |
| GO:0032012 | regulation of ARF protein signal transdu... | 6 | 1 | 0.04 | 0.03480 |
| GO:0046578 | regulation of Ras protein signal transdu... | 6 | 1 | 0.04 | 0.03480 |
| GO:0008152 | metabolic process | 1857 | 15 | 10.92 | 0.03668 |
| GO:0006807 | nitrogen compound metabolic process | 1505 | 13 | 8.85 | 0.04100 |
| GO:0006886 | intracellular protein transport | 56 | 2 | 0.33 | 0.04170 |
| GO:0034613 | cellular protein localization | 60 | 2 | 0.35 | 0.04728 |
| GO:0070727 | cellular macromolecule localization | 60 | 2 | 0.35 | 0.04728 |

Table 6: GO enrichment analysis for genes with domain rearrangements at the origin of termites in *R. speratus*

| GO.ID | Term | Annotated | Significant | Expected | pvalue |
|---|---|---|---|---|---|
| GO:0006259 | DNA metabolic process | 105 | 2 | 0.18 | 0.011 |
| GO:0005996 | monosaccharide metabolic process | 11 | 1 | 0.02 | 0.018 |

Table 7: GO enrichment analysis for genes with domain rearrangements at the origin of termites in *C. formosanus*

| GO.ID | Term | Annotated | Significant | Expected | pvalue |
|---|---|---|---|---|---|
| GO:0006281 | DNA repair | 34 | 3 | 0.3 | 0.0033 |
| GO:0006974 | cellular response to DNA damage stimulus | 34 | 3 | 0.3 | 0.0033 |
| GO:0033554 | cellular response to stress | 34 | 3 | 0.3 | 0.0033 |
| GO:0007155 | cell adhesion | 51 | 3 | 0.46 | 0.0103 |
| GO:0022610 | biological adhesion | 51 | 3 | 0.46 | 0.0103 |
| GO:0006950 | response to stress | 58 | 3 | 0.52 | 0.0145 |

Table 8: GO enrichment analysis for genes with domain rearrangements at the origin of termites in *M. natalensis*

| GO.ID | Term | Annotated | Significant | Expected | pvalue |
|---|---|---|---|---|---|
| GO:0006811 | ion transport | 228 | 9 | 2.23 | 0.00022 |
| GO:0006486 | protein glycosylation | 31 | 2 | 0.3 | 0.03621 |
| GO:0009101 | glycoprotein biosynthetic process | 31 | 2 | 0.3 | 0.03621 |
| GO:0043413 | macromolecule glycosylation | 31 | 2 | 0.3 | 0.03621 |
| GO:0070085 | glycosylation | 31 | 2 | 0.3 | 0.03621 |
| GO:0009100 | glycoprotein metabolic process | 32 | 2 | 0.31 | 0.03840 |
| GO:0030030 | cell projection organization | 5 | 1 | 0.05 | 0.04808 |
| GO:0120036 | plasma membrane bounded cell projection ... | 5 | 1 | 0.05 | 0.04808 |

Table 9: GO enrichment analysis for genes with domain rearrangements at the origin of true workers in *R. speratus*

| GO.ID | Term | Annotated | Significant | Expected | pvalue |
|---|---|---|---|---|---|
| GO:0006811 | ion transport | 226 | 14 | 2.34 | 1.1e-08 |
| GO:0055085 | transmembrane transport | 344 | 15 | 3.57 | 3.3e-07 |
| GO:0006810 | transport | 580 | 17 | 6.01 | 1.1e-05 |
| GO:0051234 | establishment of localization | 583 | 17 | 6.04 | 1.2e-05 |
| GO:0051179 | localization | 594 | 17 | 6.16 | 1.5e-05 |
| GO:0048193 | Golgi vesicle transport | 15 | 2 | 0.16 | 0.010 |
| GO:0009987 | cellular process | 2330 | 29 | 24.16 | 0.020 |
| GO:0006486 | protein glycosylation | 29 | 2 | 0.3 | 0.036 |
| GO:0009101 | glycoprotein biosynthetic process | 29 | 2 | 0.3 | 0.036 |
| GO:0043413 | macromolecule glycosylation | 29 | 2 | 0.3 | 0.036 |
| GO:0070085 | glycosylation | 29 | 2 | 0.3 | 0.036 |
| GO:0009100 | glycoprotein metabolic process | 31 | 2 | 0.32 | 0.040 |

Table 10: GO enrichment analysis for genes with domain rearrangements at the origin of true workers in *C. formosanus*

| GO.ID | Term | Annotated | Significant | Expected | pvalue |
|---|---|---|---|---|---|
| GO:0006811 | ion transport | 140 | 12 | 3.11 | 5.3e-05 |
| GO:0006313 | transposition, DNA-mediated | 13 | 4 | 0.29 | 0.00014 |
| GO:0032196 | transposition | 13 | 4 | 0.29 | 0.00014 |
| GO:0006310 | DNA recombination | 16 | 4 | 0.36 | 0.00034 |
| GO:0055085 | transmembrane transport | 240 | 14 | 5.33 | 0.00076 |
| GO:0015074 | DNA integration | 49 | 4 | 1.09 | 0.02269 |
| GO:0006810 | transport | 482 | 17 | 10.71 | 0.03565 |
| GO:0051234 | establishment of localization | 484 | 17 | 10.75 | 0.03689 |
| GO:0051179 | localization | 490 | 17 | 10.89 | 0.04081 |

Table 11: GO enrichment analysis for genes with domain rearrangements at the origin of true workers in *M. natalensis*

| Species | N of ancRA | N of non-ancRA | mean CpGo/e for ancRA | mean CpGo/e for non-ancRA | p-value |
|---|---|---|---|---|---|
| *B. germanica* | 65 | 11362 | 0.555 | 0.603 | 0.116 |
| *D. punctata* | 68 | 13751 | 0.542 | 0.543 | 0.597 |
| *P. americana* | 63 | 14782 | 0.655 | 0.615 | 0.108 |

Table 12: CpGo/e in ancestral arrangements (ancRA) in cockroaches that underwent changes in termites. P-value for one-sided Wilcoxon test

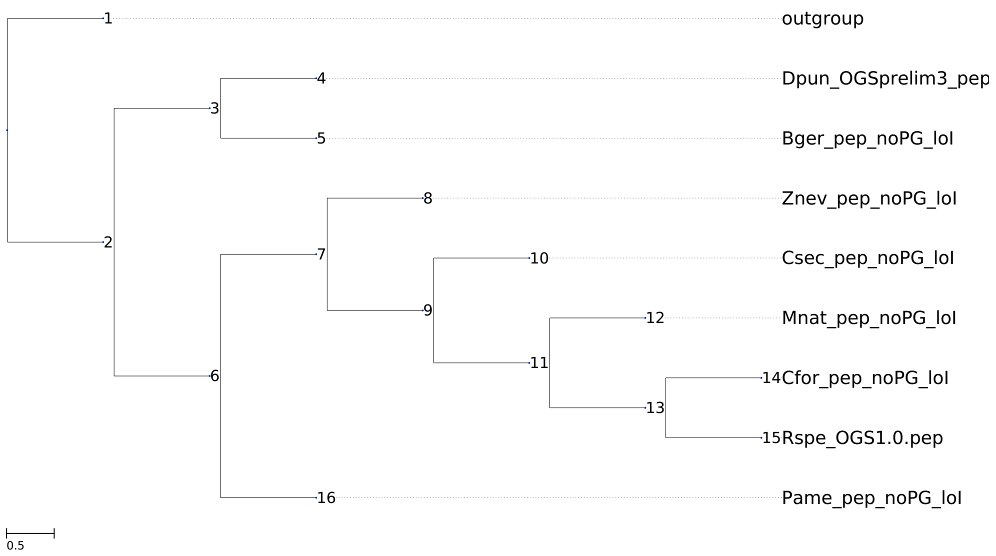


Figure 1: Tree topology used for the domain rearrangement reconstruction with node numbers

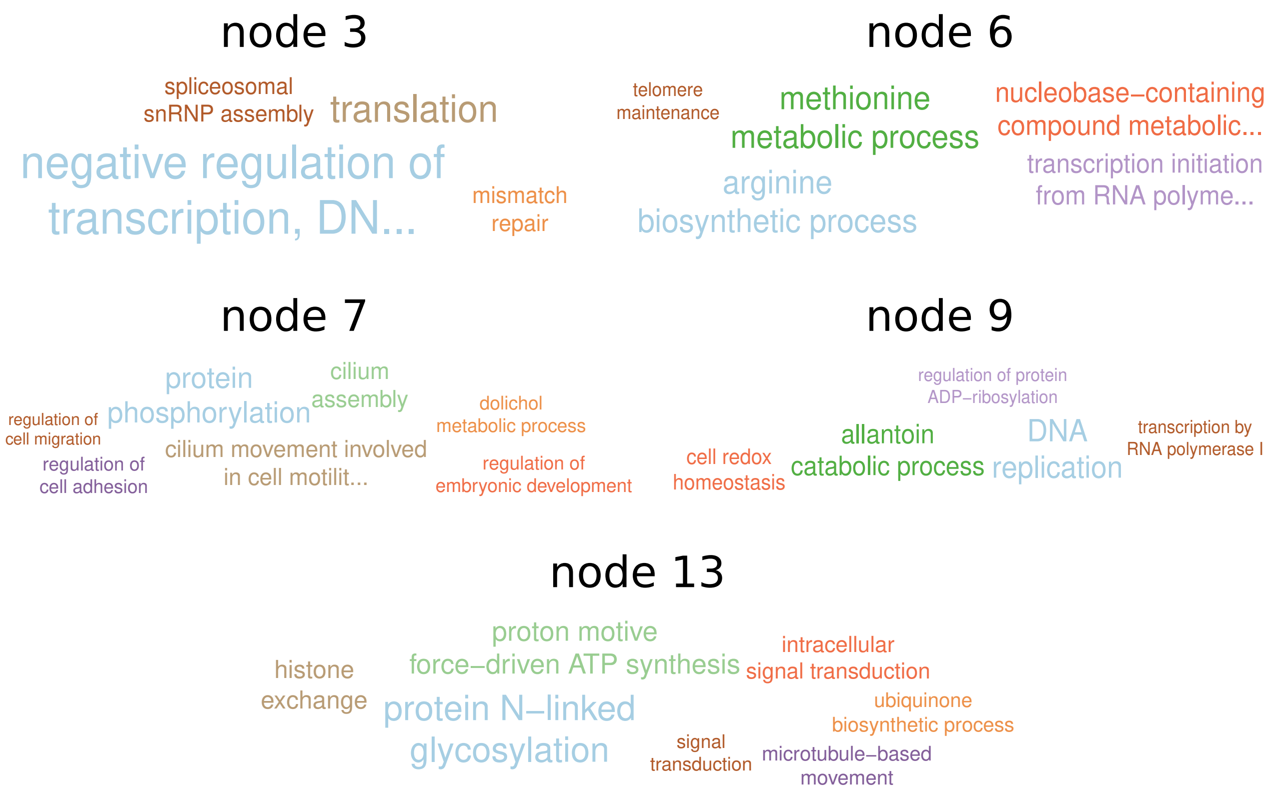


Figure 2: Enriched GO terms of rearranged domains on different nodes (node numbers from Supplementary Figure 1)

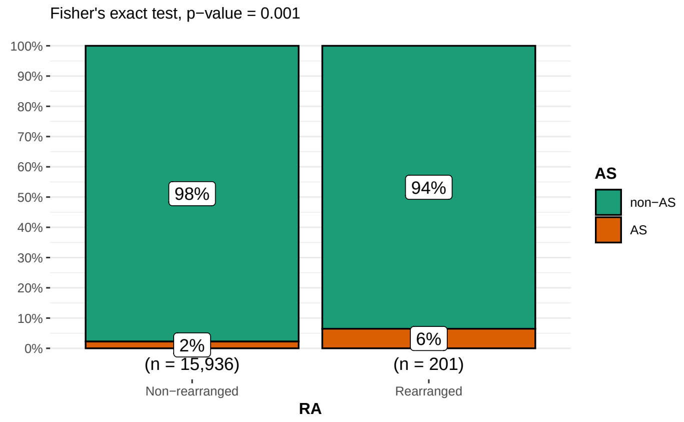


(a) Alternatively spliced in workers

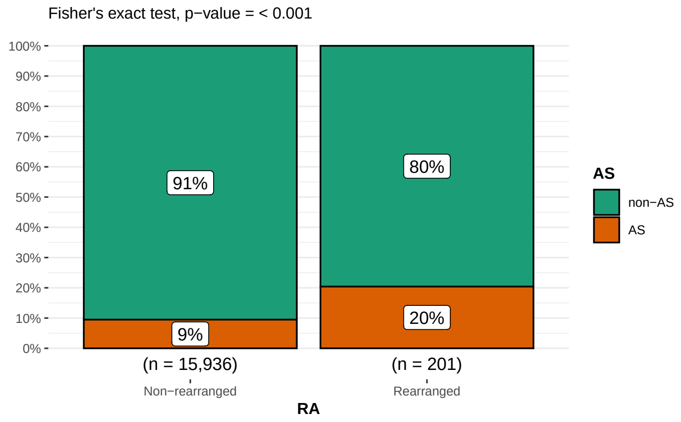


(b) Alternatively spliced in kings

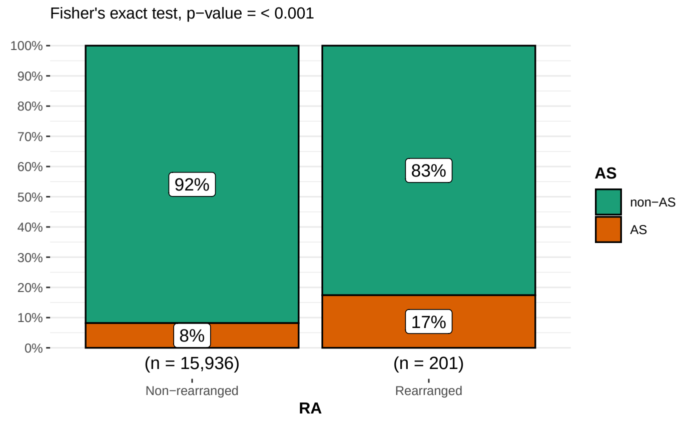


(c) Alternatively spliced in alate queens

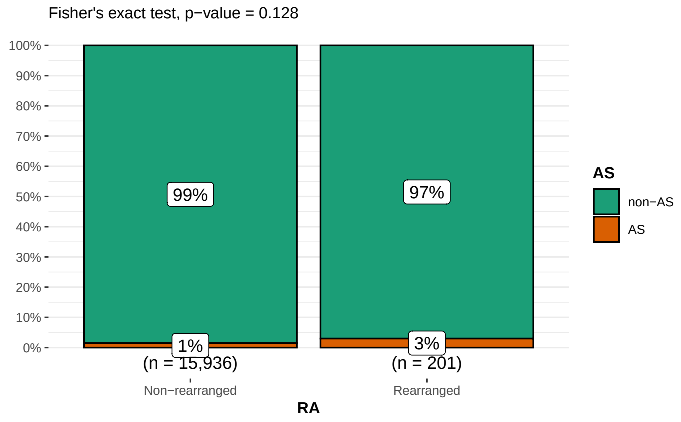


(d) Alternatively spliced in mature queens

Figure 3: The proportions of genes that are alternatively spliced (AS) in different castes within non-rearranged genes and genes rearranged in the origins of termites and true workers in *M. natalensis*

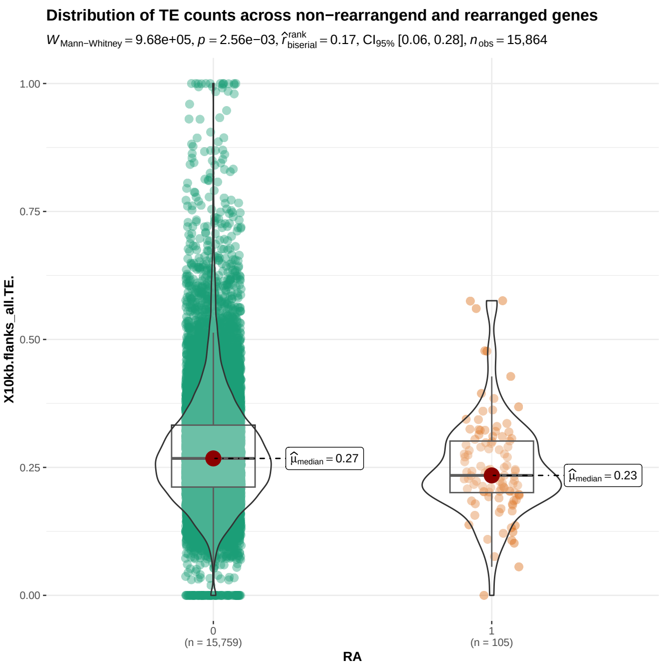


(a) *Z. nevadensis*

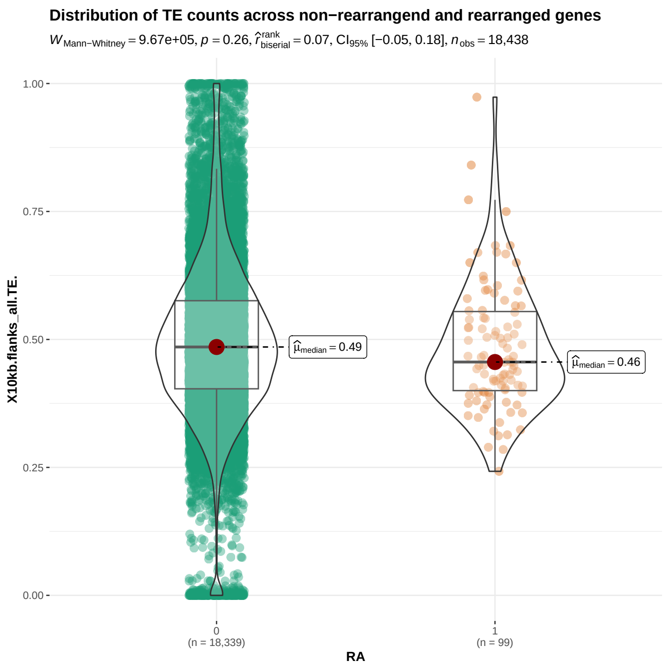


(b) *C. secundus*

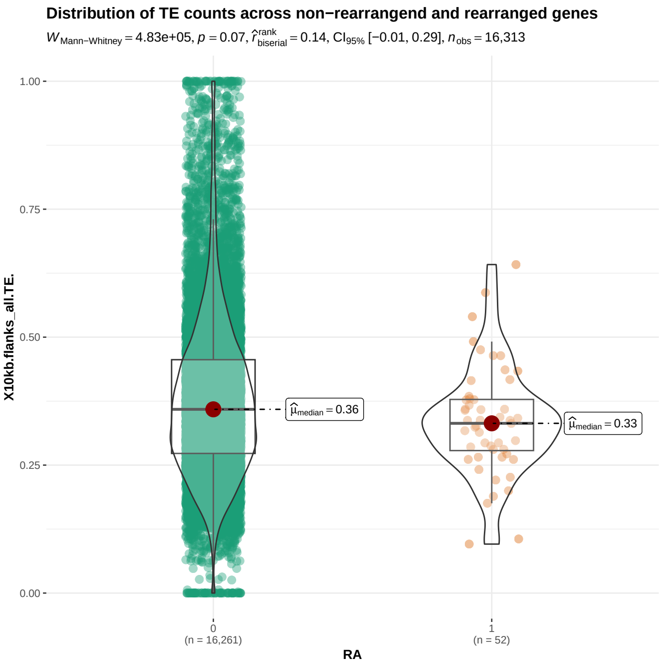


(c) *M. natalensis*, termite root

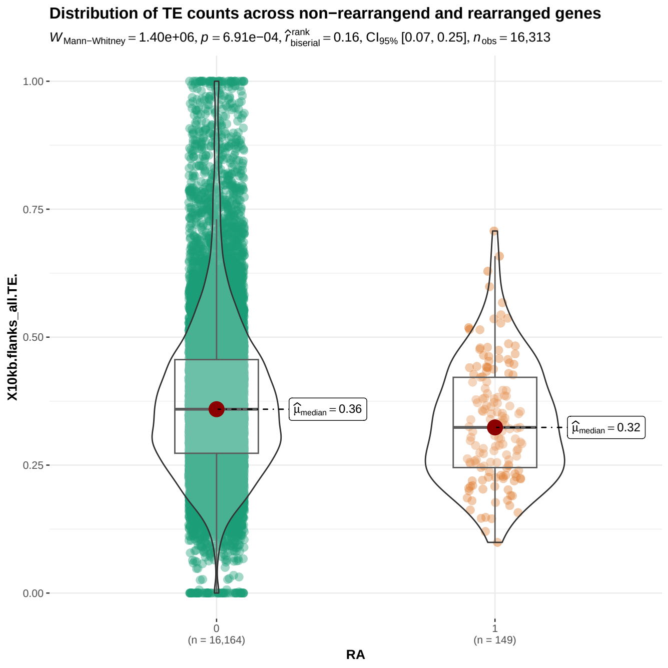


(d) *M. natalensis*, true worker root

Figure 4: TE proportions in the 10kb flanking regions of rearranged (RA: 1) and non-rearranged (RA: 0) genes in four termite species

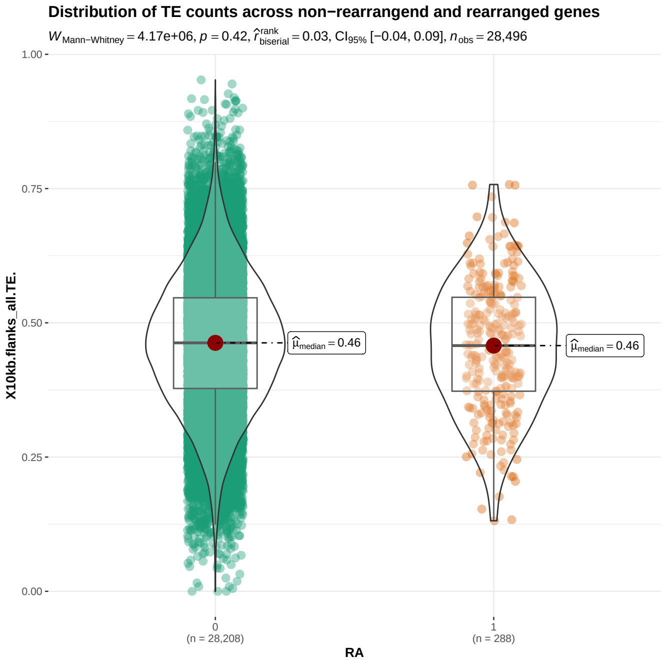


Figure 5: TE proportions in 10kb flanking regions of rearranged (RA: 1) and non-rearranged (RA: 0) genes in *B. germanica*.
